# Mechanistic Insights into G-protein Activation via Phosphorylation Mediated Non-Canonical Pathway

**DOI:** 10.1101/2024.01.15.575647

**Authors:** Kunal Shewani, Midhun K. Madhu, Rajesh K. Murarka

**Affiliations:** Department of Chemistry, Indian Institute of Science Education and Research Bhopal, Bhopal Bypass Road, Bhopal 462066, MP, India; Department of Biological Sciences, Indian Institute of Science Education and Research Bhopal, Bhopal Bypass Road, Bhopal 462066, MP, India

## Abstract

Activation of heterotrimeric G-proteins (G*αβγ*) downstream to receptor tyrosine kinases (RTKs) is a well-established crosstalk between the signaling pathways mediated by G-protein coupled receptors (GPCRs) and RTKs. While GPCR serves as a guanine exchange factor (GEF) in the canonical activation of G*α* that facilitates the exchange of GDP for GTP, the mechanism through which RTK phosphorylations induce G*α* activation remains unclear. Recent experimental studies revealed that the epidermal growth factor receptor (EGFR), a well-known RTK, phosphorylates the helical domain tyrosine residues Y154 and Y155 and accelerates the GDP release from the G*α*i3, a subtype of G*α*-protein. Using well-tempered metadynamics and extensive unbiased molecular dynamics simulations, we captured the GDP release event and identified the intermediates between bound and unbound states through Markov state models. The additional negative charges introduced by phosphorylations rewired the inter-residue interactions and significantly weakened the salt bridges at the domain interface, contributing to the increased separation of the Ras-like and helical domains of G-protein. Furthermore, the unfolding of helix *α*F resulted in greater flexibility near the hinge region, facilitating a greater distance between domains in the phosphorylated G*α*i3. The release of GDP in the phosphorylated G-protein occurred at a faster rate compared to the unphosphorylated state, caused by increased fluctuations in conserved regions of P-loop, switch 1, and switch 2. Overall, this study provides atomistic insights into the activation of G-proteins induced by RTK phosphorylations and identifies the specific structural motifs involved in the process. The knowledge gained from the study could establish a foundation for targeting non-canonical signaling pathways and developing therapeutic strategies against the ailments associated with dysregulated G-protein signaling.

## 1 Introduction

Heterotrimeric G-proteins (G*αβγ*) act as signal transducers, enabling the flow of information from the extracellular region to the intracellular downstream effector proteins that control various physiological processes.(1, 2) The process of signal transduction begins upstream, where small molecular agonists, hormones, peptides, light, and other chemical and physical stimuli are received by transmembrane receptor proteins,(3) such as G-protein coupled receptors (GPCRs),(4, 5) ion channels,(6, 7) and enzyme-linked receptors.(8, 9) G-protein-mediated signaling induced by GPCRs is implicated in almost all physiological processes, including vision, gustation, olfaction, immune system regulation, and homeostasis maintenance.(10) In GPCR signaling, the guanine diphosphate (GDP) bound G*α* subunit of the heterotrimeric G-protein attaches the activated receptor on its intracellular surface, leading to the exchange of GDP with guanine triphosphate (GTP); GPCRs function as a guanine-nucleotide exchange factor (GEF) that facilitates the nucleotide replacement.(1, 4, 5) Subsequently, the GTP-bound trimer dissociates from the receptor, followed by the separation of G*α*-GTP monomer and G*βγ* dimer, initiating their respective downstream signaling. Lastly, the inactive state of G-proteins is attained by GTP hydrolysis and recoupling of the G*α* subunit to the G*βγ* dimer.(11) In humans, there are 16 G*α* proteins classified into four families: G*α*s, G*α*i, G*α*q, and G*α*12. These G-proteins play crucial roles in regulating various cellular functions, including inositol phosphate signaling, Rho activation, cAMP production, and Ca^2+^ modulation.(12)

Experimental studies have demonstrated that the interdomain separation of the Ras-like and *α*-helical domains of the G*α* subunit (Fig. 1) results in the release of GDP upon GPCR engagement.(13–15) However, a recent study based on molecular dynamics (MD) simulations and experiments, revealed that domain separation is not sufficient and conformational rearrangements in the Ras-like domain are necessary for the release of GDP. (16) Insertion of helix-5 (*α*5) of G*α* subunit into the cytosolic cleft of a GPCR, leads to a shift of its position with respect to the rest of the Ras-like domain, resulting in allosteric changes that promote the detachment of GDP. (16) Notably, a recent computational study has shown that the rate-limiting step for GDP release is correlated with the tilt of the H5 rather than the translation.(17)

**Figure 1:**
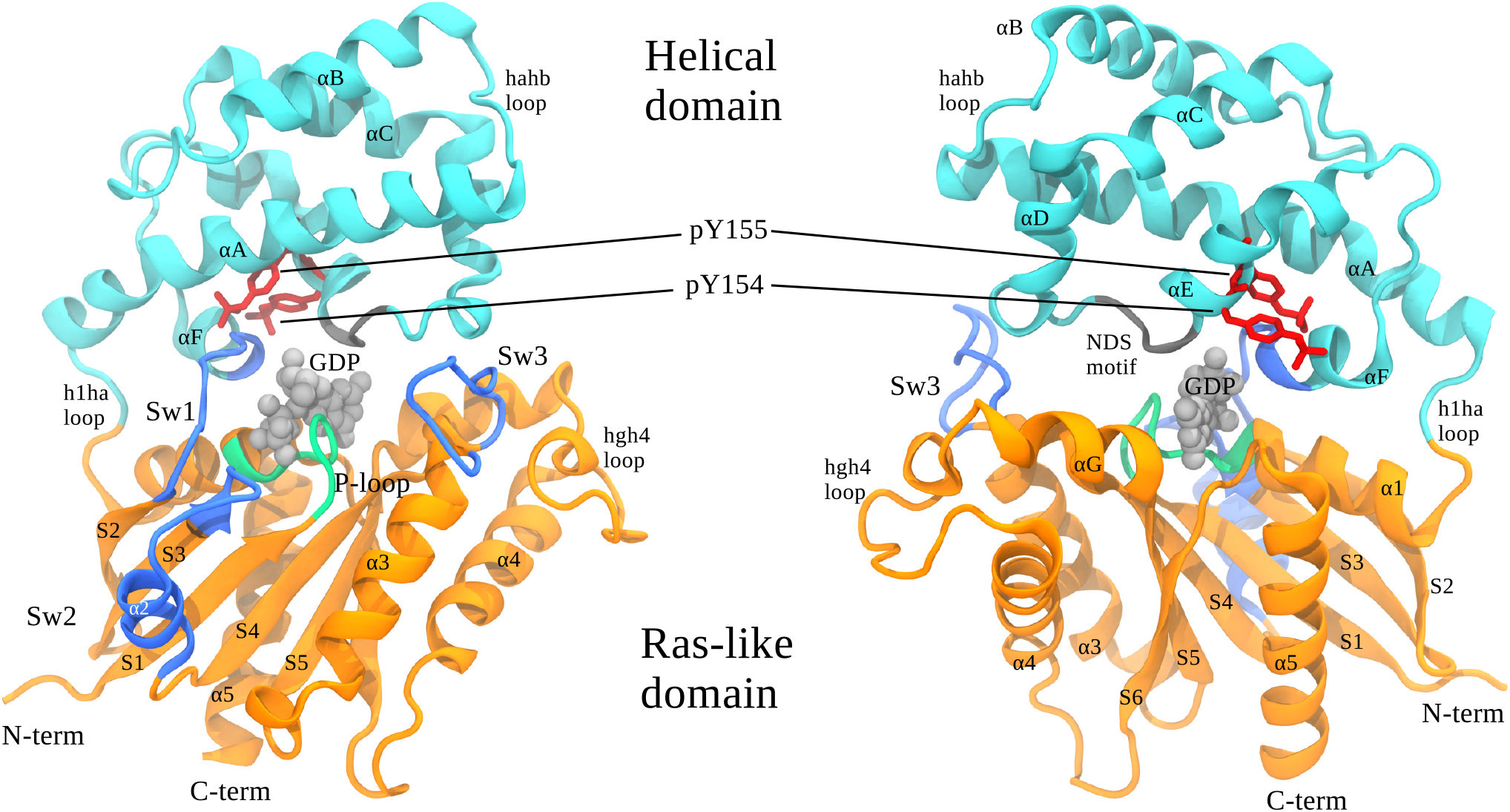
G-protein (G*α*i3) structure in cartoon representation Ras-like domain is in orange color, *α*-helical domain is in cyan color, and GDP is in space-filling representation in silver color. Conformational switches (Sw1, Sw2, and Sw3) are highlighted in blue, P-loop in green, NDS-motif in gray, and phosphorylation sites are represented in red color.

G-protein signaling is further modulated by various factors, such as guanine nucleotide dissociation inhibitors (GDIs), GTPase-activating proteins (GAPs), and other accessory proteins. (18) In addition to GPCR-GEFs, guanine nucleotide exchange modulators (GEMs) constitute a distinct class of proteins capable of independently regulating G-protein activity and acting as GEFs.(19) These non-GPCR GEFs (GEM-GEFs) share an evolutionarily conserved GEM motif (20) and functionally differ from each other. (21–25)

The GEMs with G*α*-binding and -activating (GBA) motifs, such as Girdin (G*α*-interacting vesicle-associated protein or GIV), dishevelled-associating protein with a high frequency of leucines (DAPLE), nucleobindin 1 (NUCB1 or Calnuc), and nucleobindin 2 (NUCB2), are known to activate G-proteins.(20, 22, 23) However, it is important to note that G-protein activation by GEM-GEFs differs from that of GPCR-GEFs.(26) Notably, GPCRs form major contacts with *α*5 helix of G*α* (27), while different GEMs having GBA motifs, such as DAPLE and GIV, bind to the switch 2 (sw2)/helix-3 (*α*3) region of G*α*i (Fig. 1) as demonstrated by NMR experiments. (25) Moreover, GPCRs bind to the heterotrimeric form of G-proteins, whereas GEMs mostly interact with the G*α* subunit, as GEMs and G*βγ* share a common binding interface on G*α*. A recent study reported the structure and dynamics of G*α*i3, a subtype of G*α*i, bound to GIV and identified the role of specific conserved conformational switches in GEM-mediated G-protein activation.(28)

Furthermore, a number of studies have suggested the existence of crosstalk between signaling via receptor tyrosine kinases (RTKs), another class of cell surface receptors, and GPCR signaling.(29–37) The ligand binding to RTKs, such as epidermal growth factor receptor (EGFR), leads to the autophosphorylation of their cytoplasmic tails; this allows the recruitment and phosphorylation of various adapter proteins that mediate signal transfer to the cell interior.(38–40)

It is known that G-proteins are also activated downstream of RTKs (41), and one of the mechanisms suggested is the phosphorylation of tyrosine residues in G-proteins. (42–50) A recent study using various biochemical and biophysical experiments identified the tyrosine residues (Y154 and Y155) present in the helical domain of G*α*i3 (Fig. 1) are prone to EGFR phosphorylation, thereby accelerate the GDP to GTP exchange.(51) It also demonstrated that the phosphorylation events are possible if G*α*i3 is bound to GIV (GEM-GEF), which acts as a scaffold for EGFR. Using linear ion-trap mass spectrometry combined with biochemical experiments further showed that the phosphorylation of serine/threonine/tyrosine residues in various regions impacts the dynamics of G*α*i3, with phosphorylation of Y320 (in Ras-like domain) separates the canonical (GPCR-dependent) G-protein signaling from the non-canonical one.(52)

Despite these experimental findings, the molecular mechanism by which tyrosine phosphorylations in G*α*i3 accelerate the nucleotide exchange remains elusive. Particularly, it is intriguing to know how the phosphorylation sites (Y154 and Y155) that are present in the helical domain influence the interactions of GDP with the Ras-like domain, leading to its dissociation. To develop a detailed understanding of the GDP dissociation process, we performed well-tempered metadynamics and conventional atomistic molecular dynamics simulations (aggregated simulation time of > 60 *μ*s) for the phosphorylated as well as unphosphorylated G*α*i3 systems. By employing the Markov state model (MSM) on the simulated data, we characterized the metastable states sampled in the process. Further, machine learning classifiers were utilized to identify important protein structural units associated with the release of the nucleotide.

The rest of the article is organized as follows. Section II describes the specifics of modeling the systems and the details of molecular dynamics simulations. Additionally, a brief outline of the string method, and Markov state model are provided in Sec. II. Section III presents and discusses the results, and finally, section IV highlights the concluding remarks of the study.

## 2 Methods

### 2.1 Modeling and system preparation

Due to the unavailability of GDP-bound inactive heterotrimeric G*α*i3 structure, the initial configuration of GDP bound G*α*i3 (residue A30^G.hns1.1^ to N347^G.H5.19^) was modeled from the crystallographic structure PDB ID: 6MHF(28) after removing the bound GIV molecule. In this context, the superscript on residues follows a common G*α* numbering scheme.(53) For instance, the notation K46^G.H1.1^ denotes that Lys46 belongs to the Ras-like domain (commonly known as GTPase domain or G) and is the first residue of helix-1 (*α*1). Similarly, the residues in the helical domain are denoted by the superscript H. Further, hns1 designates the loop between helix-N (*α*N) and *β* strand-1 (S1). The model was then phosphorylated at the residues Y154^H.HE.4^ and Y155^H.HE.5^, and thereafter, both the phosphorylated and the unphosphorylated systems were solvated using CHARMM-GUI web server(54–56) with explicit water molecules utilizing the CHARMM-modified TIP3P water model.(57) To achieve natural ionic concentration of 150mM, potassium (K^+^) and chloride (Cl^-^) ions were added to the simulation box. The modeled proteins were capped at the N-terminus with an acetyl group and the C-terminus with a methylamine group. The forcefield used for protein was CHARMM36m,(58) and the parameters for GDP were obtained from the ParamChem server using CHARMM General Force Field (CGenFF).(59) The unphosphorylated G*α*i3 system contained 54961 atoms, with 16570 water molecules, 53 K^+^ ions, and 47 Cl^-^ ions, whereas the phosphorylated G*α*i3 system contained 54905 atoms, with 16548 water molecules, 57 K^+^ ions, and 47 Cl^-^ ions. The initial volume of both systems was approximately measured to be 97×87×73 Å^3^.

### 2.2 Molecular dynamics simulations

We used GROMACS version 2019.5 for conventional molecular dynamics (cMD) simulations.(60–66) Initially, both the systems were energy minimized for 5000 steps using the steepest descent algorithm. The temperature of the systems was gradually raised from 0 K to 310 K for the duration of 125 ps in the NVT ensemble, where restraints were applied on protein-heavy atoms. Subsequently, a 100 ns NPT equilibration without any restraints was carried out before running production runs. The production simulations were performed under the NPT ensemble at 310 K temperature using a Nose-Hoover(67) thermostat with a 1 ps coupling constant. The pressure was maintained at 1 bar using a Parrinello-Rahman(68) barostat with a 5 ps coupling constant. Long-range electrostatic interactions were calculated using the particle mesh Ewald method (PME)(69) with a grid spacing of 1 Å. The real space of the electrostatic interactions and the Lennard-Jones potential for the van der Waals interactions were evaluated with a cut-off of 12 Å, and a force switch was applied at 10 Å. The LINCS(70) algorithm was used to constrain the bonds to hydrogen atoms. A timestep of 4 fs was used in simulations by employing hydrogen mass repartitioning.(71)

### 2.3 Well-tempered metadynamics

The biomolecular processes of interest generally happen at a very long time scale, which is difficult to achieve using cMD simulations; thus, enhanced sampling techniques were developed to overcome this issue. Well-tempered-metadynamics (WTMetaD), a widely used enhanced sampling technique, was employed in this study to achieve GDP dissociation from G-protein. In classical metadynamics,(72) a history-dependent bias potential is introduced as a function of time and a set of predefined collective variables (CVs) that are functions of the positions of particles in the system, along which the evolution of the system can be observed. The bias potential is in the form of a Gaussian defined as,

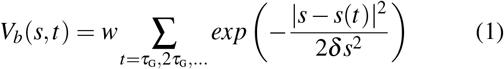

where *s* is a given collective variable, *δs* is the width, *w* is the height, and *τ*_*G*_ is the time interval at which the Gaussian is deposited. This bias potential (*V*_*b*_) allows the system to escape local minima by filling up the wells in the free energy surface of the CVs. After a certain amount of period, this deposited bias potential can be used to estimate the underlying free energy surface of the process along the relevant CV, where it can be calculated as,

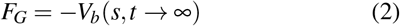

Since the height of the deposited Gaussian potential is fixed, this method has several limitations, such as overestimation of the free energy and slow convergence, as the system can explore the regions of configurational space that are physically not relevant. To alleviate these limitations, we employ a variant of metadynamics known as well-tempered metadynamics.(73, 74) In this method, the height of the Gaussian is rescaled based on an adaptive bias using the following relationship,

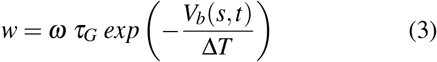

where *ω* is the bias deposition rate with the unit of energy per unit time, Δ*T* is the tunable parameter with the dimension of temperature (adjust the height of the Gaussian), and *V*_*b*_(*s, t*) is the estimate of the bias potential at the current CV position and time. In WTMetaD, the deposited Gaussian bias potential is used to approximate the free energy surface as,

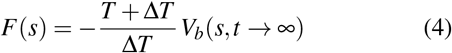

where, Δ*T* is given by the bias factor 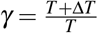. Initially, in WTMetaD, the height of the Gaussian bias potential is large, allowing it to fill the energy basins quickly. As the phase space exploration progresses, the height is scaled, which allows the free energy surface of the system to converge smoothly.

The WTMetaD simulations were performed using GRO-MACS version 2019.5, patched with plumed version 2.6.0.(75) The set of CVs used in WTMetaD simulations were defined as follows: CV1 was the distance between the center of mass of the backbone atoms of K46^G.H1.1^-T48^G.H1.3^ and phosphate atoms of GDP, and CV2 was the RMSD of GDP to the initial bound structure. The same set of CVs was also considered in a recent simulation study of G-protein.(17) Three independent simulations of 200 ns were carried out for unphosphorylated and phosphorylated G-proteins. The initial height of the Gaussian bias potential employed was set at 1.5 kJ/mol, with a width of 0.1 Åfor both the CVs. The bias factor was chosen to be 10, and the Gaussian was deposited every 2 ps.

### 2.4 String method

The free energy surface thus obtained for the GDP dissociation process was used to identify the minimum energy path (MEP) using the string method.(76) In this method, metastable states are localized around the minima of the potential *V* (*q*). Assuming *V* (*q*) has at least two minima *A* and *B*, one look for minimal energy paths (MEPs) connecting these states. By definition, MEP is a smooth curve *φ*^*^ connecting *A* and *B* which satisfies

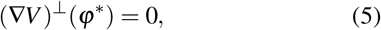

where (∇*V*)^⊥^ is the component of ∇*V* normal to *φ*^*^. Let *φ* be a string (not necessarily an MEP but an initial guess) connecting *A* and *B*. A simple method to find the MEP is to evolve *φ* according to

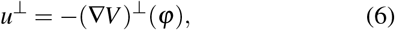

where *u*^⊥^ denotes the normal velocity of *φ*, since stationary solutions of eq. (6) satisfies eq. (5). For numerical convenience, eq. (6) can be rewritten with a parameterization term. The instantaneous position of the string is denoted as *φ*(*α, t*), where *α* is some suitable parameterization, using which eq. (6) can be written as,

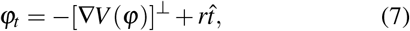

where 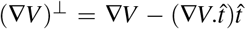, and 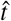 is the unit tangent vector along *φ*, 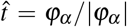. The scalar field *r ≡ r*(*α, t*) is a Lagrange multiplier uniquely determined by choice of parametrization. The simplest example is to parametrize *φ* by arc length normalized so that *α* = 0 at *A, α* = 1 at *B*. Then eq. (7) must be supplemented by the constraint (|*φ*_*α*_ |)_*α*_ = 0, which determines *r*.

Herein, after determining the MEP, 50 equally spaced conformations along the dissociation pathway were selected for each of the three WTMetaD runs as starting points for unbiased parallel MD simulations. We performed 150 independent cMD runs of 200 ns each for both unphosphorylated and phosphorylated systems, resulting in the aggregate of a total 60 *μs* data.

### 2.5 Markov state model (MSM)

The construction of a Markov State Model (MSM) serves as a potent tool for the automatic identification of relevant metastable states and the transition rates between them based on simulated trajectories.(77–82) We used PyEMMA(83) to create and analyze the MSMs derived from a cumulative 30 *μ*s of MD simulations for each system. To define the input features, we employed heavy-atom contacts between GDP and protein residues, identifying all pairs that were in 6 Å cutoff distance from the GDP molecule. To reduce the high dimensionality of the input data, we applied a linear transformation technique, known as the time-structured Independent Component Analysis (tICA). (84–86) The correlation lag time of 20 ns was chosen based on systematic construction of tICA with varied lag times, exhibiting a converged tIC surface along the first two dominant modes (Fig. S1). We considered a kinetic variance (measured as 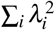, where *λ*_*i*_ represents the eigenvalue of the *i*^th^ mode)(87) of 95% to determine the number of tIC dimensions for projection of the simulated data. We employed the k-means clustering algorithm (88) to discretize the projected data into 100 clusters, chosen using VAMP2 score.(89) Subsequently, 100 microstate MSMs were constructed at varying lag times to identify the most suitable one. A lag time of 10 ns was considered for the final MSM construction. At this lag time the implied time scale plot reaches a plateau, ensuring the Markovian property of the model (Fig. S2). The hidden Markov model (HMM)(90) was then used to construct a coarse-grained kinetic model for each system, considering the number of metastable states that accounted for a separation of slower processes from the faster ones (Fig. S2). We obtained three and four-state models for the unphosphorylated and phosphorylated systems, respectively. Finally, the Chapman-Kolmogorov test (CK-test)(81) was performed that verified the Markovianity of the obtained models (Fig. S3).

### 2.6 MD simulation analysis and visualization

Principal component analysis (PCA) is a widely used dimensionality reduction technique that extracts the information about dominant motions from large dataset such as MD simulation trajectories. In this study, PCA was conducted using the GROMACS modules covar, and the anaeig module was employed to project the obtained principal components (PCs) onto the trajectories. The PCA was performed on C_*α*_ atoms of switch 2 after the protein was aligned using Ras-like domain. The root-mean-square deviations (RMSDs) and pair-wise contacts of GDP with the G-protein were computed using the built-in modules of GROMACS.(65) Distances between different groups of atoms were calculated using MDAnalysis, a Python-based trajectory analysis tool.(91) Plots were generated using the Python library Matplotlib. Visual Molecular Dynamics (VMD 1.9.4a31) was used for visualization and image rendering.(92) The non-bonding energies for the residue pairs (X and Y) were calculated as the sum of their electrostatic and van der Waals interactions 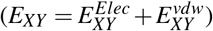 using the CPPTRAJ module of AMBER18.(93)

## 3 Results and discussion

### 3.1 WTMetaD Simulations Capture GDP Dissociation

To understand the molecular basis of the dissociation mechanism of GDP due to the phosphorylation of Y154^H.HE.4^ and Y155^H.HE.5^ in G*α*i3, we simulated the unphosphorylated and phosphorylated G-protein systems. The unbinding of nucleotide from G-protein is a long-timescale event.(16) Therefore, to accelerate and capture the process in an accessible computational time, we used well-tempered meta-dynamics (WTMetaD), a collective variable (CV)-based enhanced sampling technique (see Sec. 2.3 for details). Three independent WTMetaD simulations were performed for each system, and all of them were observed to successfully capture the dissociation of GDP, as indicated by the GDP distance from the G-protein (CV1) and GDP’s RMSD to the initial bound state (CV2) (Figs. S4a-S4d). A comparison of the average GDP contacts (Fig. S4e) revealed that the unphosphorylated G*α*i3 remained in the bound state (contact fraction ≥ 0.7) for an extended time period compared to the phosphorylated G*α*i3, whereas, the opposite was observed for the GDP dissociated state (contact fraction *≤* 0.2). This suggests that GDP dissociation from G*α*i3 is facilitated upon phosphorylation at residues Y154^H.HE.4^ and Y155^H.HE.5^.

To identify the minimum free energy pathway of GDP dissociation from G*α*i3, we utilized the string method (see Sec. 2.4) on the free energy surface (FES) obtained from independent WTMetaD simulations for each system. Subsequently, we chose conformations along the release pathway on the FES and used them as the initial seeding points for parallel unbiased simulations, generating an aggregate of 60 *μs* trajectories (see Sec. 2.4). The simulated data thus generated were used for further analysis, as discussed in the following sections.

### 3.2 Phosphorylated G-protein Shows Increased Interdomain Dynamics

The unbiased trajectories of both phosphorylated and unphosphorylated systems displayed large distances of GDP from the binding pocket (Fig. 2a), signifying the occurrence of GDP dissociation events. We examined the difference in the overall dynamics of G*α*i3 systems during the dissociation of GDP by calculating the root mean square deviation (RMSD) with respect to their initial conformations. Fig. 2b shows that the phosphorylated system has a broader distribution of RMSD values as compared to the unphosphorylated system, indicating an increased dynamics of G-protein upon phosphorylations. Similarly, an enhanced root mean square fluctuation (RMSF) of each residue was observed for the phosphorylated G*α*i3 (Fig. S5a). However, the difference in RMSDs of individual domains was relatively smaller when measured after aligning each one separately (Figs. S5b and S5c). This suggests that the increased dynamics observed for the phosphorylated G-protein are majorly influenced by the dynamics at the domain interface; in fact, the large RMSD values were found to correlate well with the interdomain separation (between Ras-like and helical domains, Fig. S5d).

**Figure 2:**
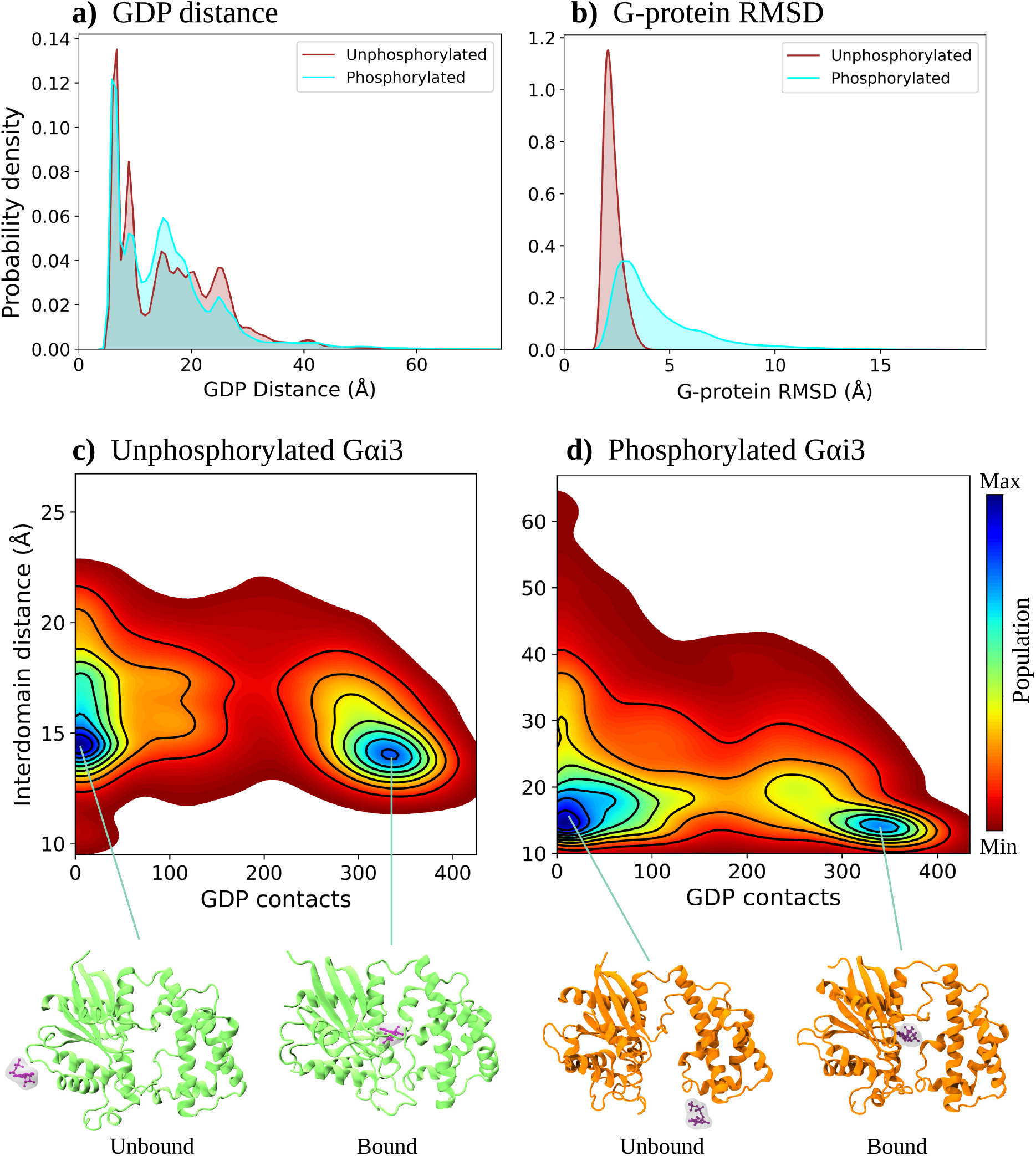
Distribution plot of unphosphorylated and phosphorylated G*α*i3 (a) Center of mass distance between GDP phosphate atoms and backbone atoms of residues K46^G.H1.1^, S47^G.H1.2^, and T48^G.H1.3^, (b) RMSD of G-protein with respect to the initial conformation. 2D density plot along GDP contacts with G-protein binding residues and interdomain distance between Ras-like and helical domains (heavy atom COM distance of residue R144^H.HD.11^ and N241^G.s4h3.15^) (c) unphosphorylated and (d) phosphorylated G*α*i3.

To obtain a comprehensive understanding of interdomain dynamics during GDP dissociation, we generated a 2D density plot correlating GDP contacts and interdomain distance for the entire data set (Figs. 2c and 2d). In the unphosphorylated G*α*i3, the interdomain distance remains below 25 Å, with an evident domain separation observed even in the bound state (Fig. 2c). In contrast, a substantially large domain opening was observed for both the GDP-bound and unbound states of phosphorylated G*α*i3 (Fig. 2d). This suggests that the presence of phosphorylated residues (pY154^H.HE.4^ and pY155^H.HE.5^) located at helix *α*E (Fig. 1) caused an increase in the extent and frequency of domain opening during the GDP dissociation. It is to be noted that the separation of G-protein domains is necessary for the release of the GDP molecule. However, previous simulation and experimental studies have indicated that domain separation alone is not sufficient for GDP dissociation; the internal structural rearrangements in the Ras-like domain are crucial for weakening the affinity of the nucleotide, (16) as will be discussed in the following section.

To find out the molecular basis of the large interdomain dynamics in the phosphorylated system, we next examined the interactions of residues located at the domain interface. The phosphorylation of Y154^H.HE.4^ and Y155^H.HE.5^ had a significant impact on the interaction energy of residue pairs involving charged amino acids, leading to the reduced stabilization (Fig. 3a). For instance, the average interaction energy (electrostatic + van der Waals) of the residues lysine K54^G.H1.9^ and glutamate E65^H.HA.3^ (separated by linker 1 connecting helical and Ras-like domains, Fig. 3a) reduced from -91.2 *kcal/mol* to -57.6 *kcal/mol* (Fig. 3a). A similar decrease in interaction energy was observed in the phosphorylated G*α*i3 for many of the interfacial residue pairs (Table S2; Fig. 3a), including R144^H.HD.11^-D229^G.s4h3.3^ and R144^H.HD.11^-D231^G.s4h3.5^ (lower region of the domain interface), and D150^H.hdhe.5^-K270^G.s5hg.1^ (central region, close to the nucleotide). We found that the reduced stability was caused due to the perturbations introduced by the negatively charged phosphate groups and their interactions with the interfacial charged residues (Table S2). In fact, a larger domain separation observed upon phosphorylation can be attributed to the weakening of these stabilizing interactions.

**Figure 3:**
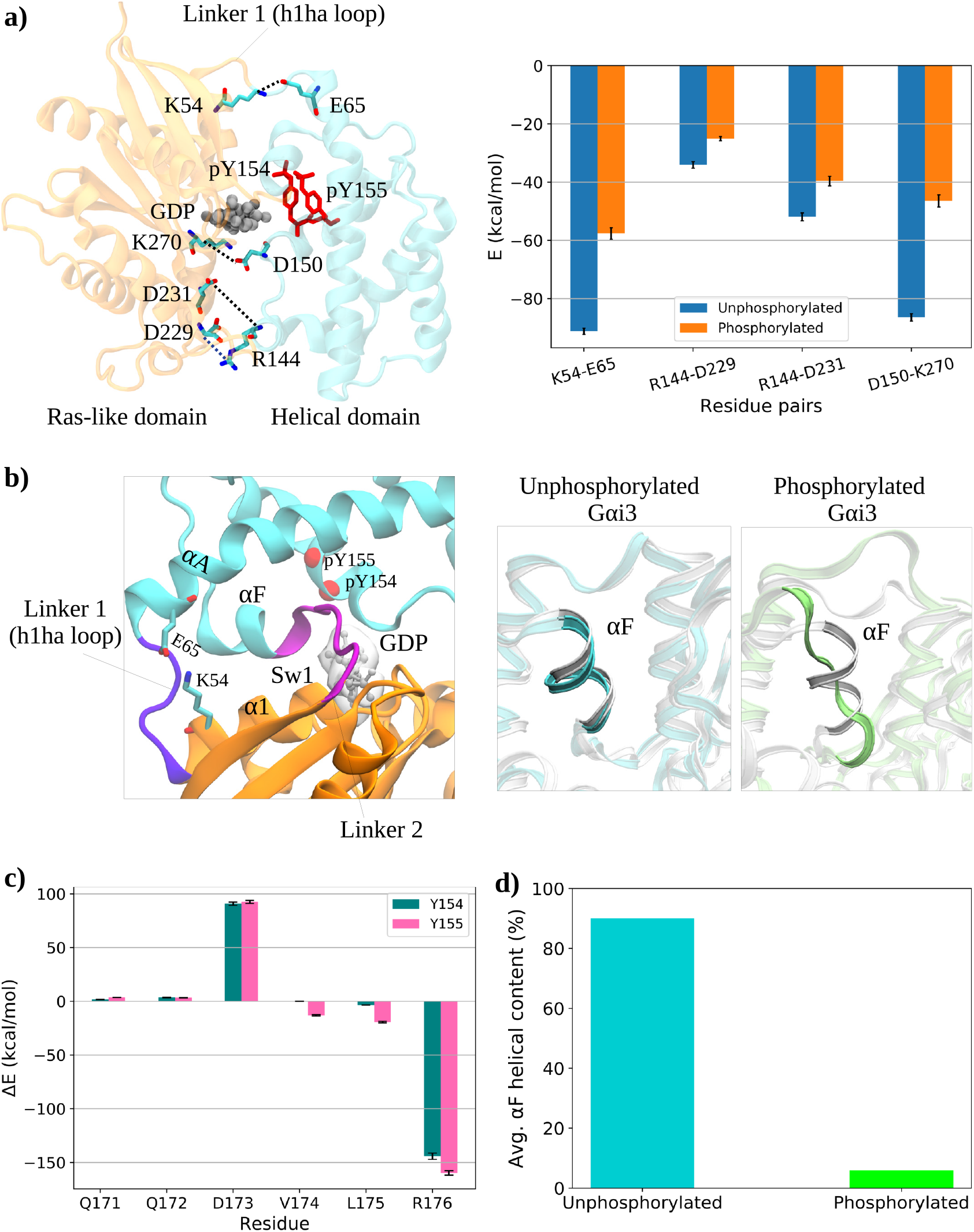
(a) Interactions of residue pairs at the domain interface and comparison between their average interaction energy of the unphosphorylated and phosphorylated systems. (b) Structure of phosphorylated G-protein with helix *α*F, linker 1 (h1ha loop), and linker 2 (sw1). The representative conformations of helix *α*F from the simulations are shown for the unphosphorylated (cyan) and phosphorylated (lime) systems overlapped with the crystal structure (light gray) of G*α*i3 (PDB ID: 6MHF). (c) Difference in average interaction energy between the *α*F residues and phosphorylation sites in the unphosphorylated and phosphorylated systems. (d) The percentage of helical content in *α*F helix during simulations. The error bars for interaction energies were obtained from the standard error of the mean (SEM).

Further, we found that the helix *α*F, which is located opposite to the *α*E and connected to the linker 2/switch 1 (sw1), unfolded in the case of phosphorylated G*α*i3 (Fig. 3b and Fig. S6). To find out the underlying reason, we calculated the difference in average interaction energy of *α*F residues with the phosphorylation sites between G*α*i3 systems. The repulsive interactions of the residues Q171^H.HF.1^-D173^H.HF.3^ (in particular, a highly unfavorable one for D173^H.HF.3^), combined with the stabilizing interactions of residues V174^H.HF.4^-R176^H.HF.6^ in the other half of *α*F (notably, a highly stablizing one for R176^H.HF.6^) with the phosphorylated pY154^H.HE.4^ and pY155^H.HE.5^ (Fig. 3c) introduces a strain in *α*F helix, thereby causing its unfolding (Fig. 3d). In effect, this large conformational change in *α*F increases the flexibility of the interdomain linker 2, which facilitates a larger separation of Ras-like and helical domains. Collectively, the destabilization of charged residue pairs at the domain interface and unfolding of the *α*F helix due to perturbations induced by the phosphorylations appear to be the underlying reasons for significant domain opening observed in the phosphorylated G*α*i3.

### 3.3 Enhanced Fluctuations in P-loop and Switch Regions Upon Phosphorylations Reduces the Affinity of GDP

It is known that both GPCRs and non-GPCR GEFs facilitate the exchange of GDP through allosterically influencing the nucleotide-binding regions in G*α*-subunit; GPCRs induce changes in P-loop and sw1,(14, 53, 94) whereas non-GPCR GEFs such as GIV perturb sw1.(25, 28) Moreover, as discussed earlier, a recent study using cellular and biochemical experiments revealed that the phosphorylation of tyrosine residues (Y154^H.HE.4^ and Y155^H.HE.5^) located in the interdomain cleft of G*α*i3 also induce enhanced G-protein activation.(51) We investigated the alterations in conserved structural elements, including the P-loop and switch regions, to understand the basis of phosphorylation-driven GDP release.

Both the P-loop and sw1 regions displayed larger RMSD values for the phosphorylated G-protein compared to the unphosphorylated one (Fig. 4a and 4b, top panel). To this effect, a closer look at the change (between the G*α*i3 systems) in the average interaction energy of residues from these structural regions with the phosphorylation sites can provide a basis for the enhanced fluctuations. The observed structural variations in the P-loop region can, in fact, be attributed to the large unfavorable and favorable interactions of residues E43^G.s1h1.4^ and K46^G.H1.1^, respectively, with the phosphorylated pY154^H.HE.4^ and pY155^H.HE.5^ (Fig. 4a, middle). Whereas for the sw1, significant stabilization of electrostatic interactions involving positively charged residues R176^H.HF.6^, R178^G.hfs2.2^, and K180^G.hfs2.4^ with the phosphorylated tyrosines, contributed to its greater structural variations (Fig. 4b, middle). Additionally, the dynamics of sw1 are also affected by its connecting helix *α*F, which was observed to undergo unfolding in phosphorylated G*α*i3 (Fig. 3b and Fig. S6). As the P-loop and sw1 regions make crucial contacts with the nucleotide, the enhanced fluctuations observed in these regions are therefore suggestive of the weakening of GDP contacts upon phosphorylation.

**Figure 4:**
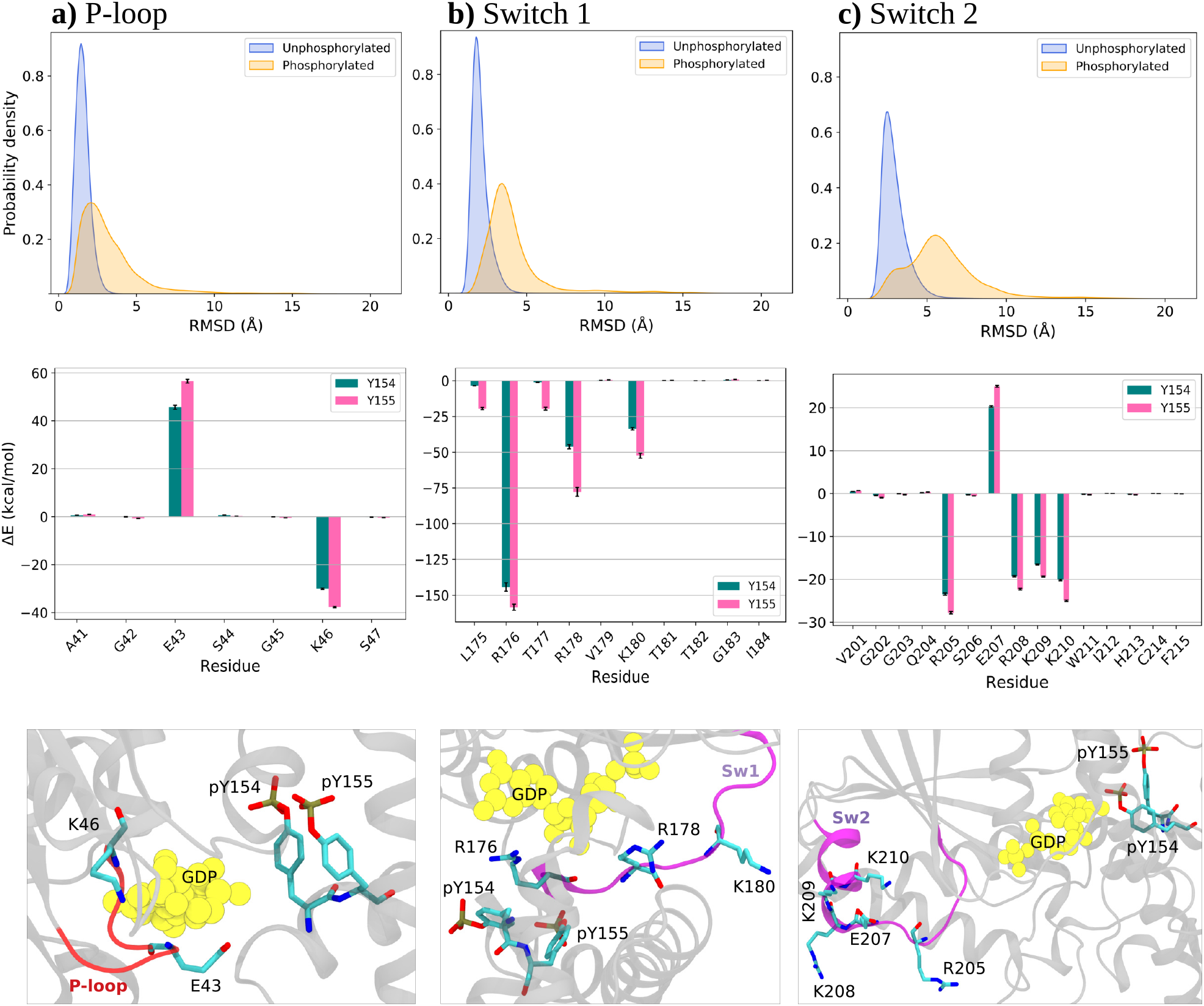
Fluctuations in conserved regions contributing towards GDP release. The top panel shows the distribution of RMSD values between unphosphorylated and phosphorylated systems, the middle panel shows the difference in average interaction energy of residues with phosphorylation sites Y154 and Y155 between the systems, and the error bars were obtained by SEM. The bottom panel shows residues of the conserved regions that have a significant change in the interaction energies for (a) P-loop, (b) switch 1, and (c) switch 2.

We further observed large fluctuations in the sw2 region of phosphorylated G*α*i3 (Fig. 4c, top) due to changes in the electrostatic interactions of several charged residues (R205^G.H2.1^, E207^G.H2.3^, R208^G.H2.4^, K209^G.H2.5^, and K210^G.H2.6^) with pY154^H.HE.4^ and pY155^H.HE.5^ (Fig. 4c, middle). In addition to the P-loop and sw1, the conformational change in sw2 is also reported to be crucial for G-protein activation, as it releases spatial constraints from the nearby structural units and accelerates GDP release from the pocket.(25, 28, 95, 96) We investigated sw2 motions by conducting principal component analysis (PCA) on its C_*α*_ atoms after aligning the Ras-like domain. The first two principal components (PCs) collectively accounted for 61.2% and 75.9% of the overall motions in unphosphorylated and phosphorylated G*α*i3, respectively. The free energy surfaces exhibit a single deep minimum for the unphosphorylated system (Fig. S7). In contrast, phosphorylated G*α*i3 displays a deep minimum with a nearby shallow one, indicating conformational heterogeneity in the sw2 region due to phosphorylation (Fig. S7). We note that a previous experimental study using mutations and mass spectrometry revealed that an increased dynamics of sw2 is correlated with a higher nucleotide exchange rate in G*α*i3.(28) Overall, the conformational and interaction energy analyses suggested an increase in fluctuations of key conserved regions of G*α*i3 upon phosphorylation that results in an accelerated GDP release.

### 3.4 Phosphorylations Induces Faster Nucleotide Release from G-protein

Next, we investigated the metastable conformational states sampled by the unphosphorylated and phosphorylated G-proteins by employing the Markov state model (MSM), a widely used method that provides thermodynamic and kinetic information of the process of interest from simulation trajectories.(82, 83) For each system, a cumulative 30 *μ*s of trajectory data were used to construct the MSM. For dimensionality reduction using the tICA method, the contacts between the heavy atoms of GDP and G-protein residues were considered as the input features (Fig. 5a). The reduced tICA space thus obtained was discretized into 100 clusters using the k-means algorithm. Subsequently, by employing a hidden Markov model (HMM), the unphosphorylated and phosphorylated systems were further coarsegrained into three and four metastable states, respectively (see Sec. 2.5 for details). The three states obtained for the unphosphorylated G*α*i3 are characterized as follows: bound state (UB); intermediate state (UI) with ∼40% reduction in average GDP contacts compared to UB; and unbound (U0) state (Fig. 5b). Whereas the four states in the phosphorylated system include the bound (PB), unbound (P0), and intermediate-1 (PI1) and intermediate-2 (PI2) with a ∼10% and ∼40% decline, respectively, in average GDP contacts with respect to the bound state (Fig. 5d).

**Figure 5:**
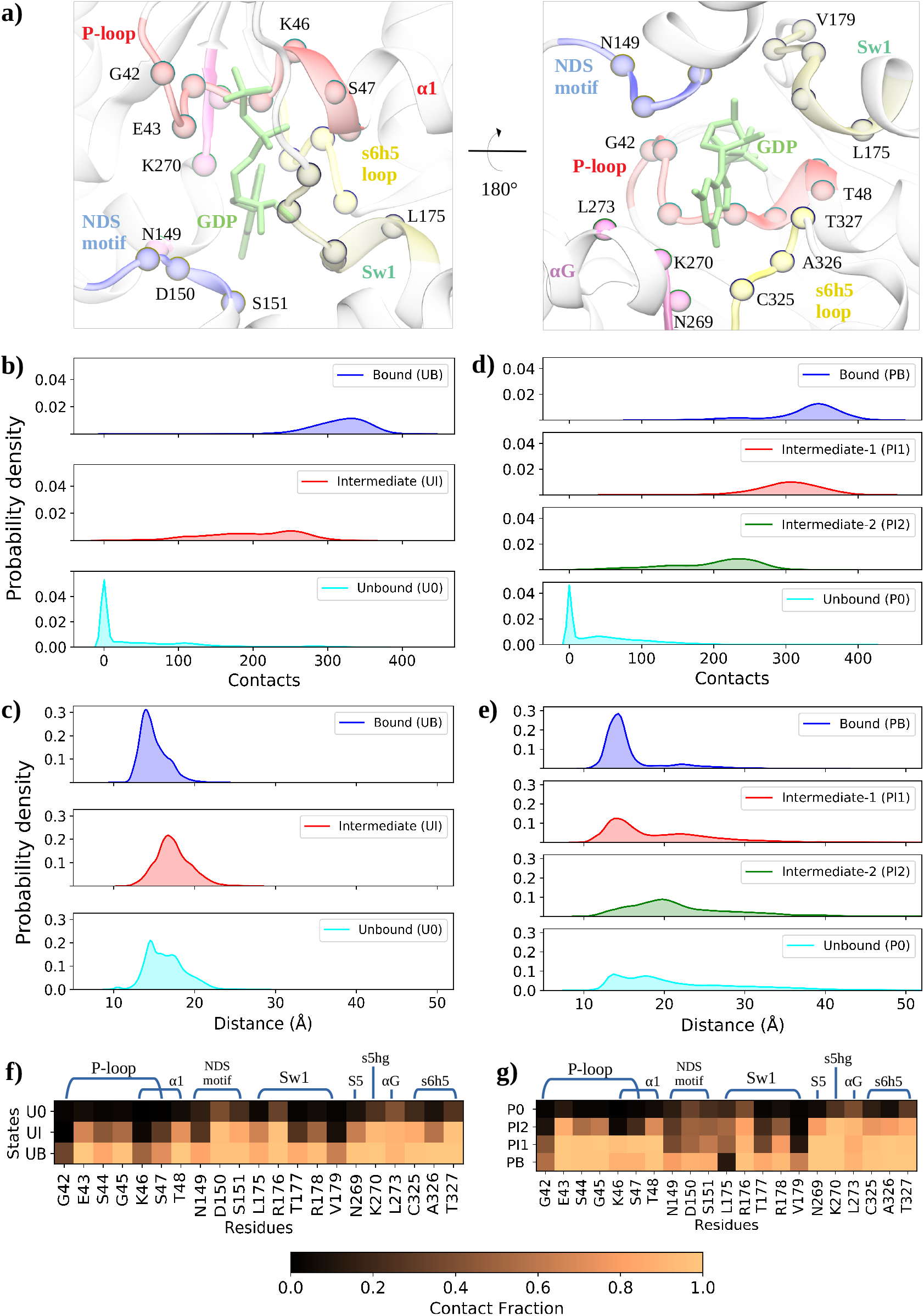
(a) Representative of GDP (lime) and its contact residues (spheres) from crystal of G-protein (PDB ID: 6MHF). Distribution plots for individual macrostates obtained from HMM. Contacts between the heavy atoms of GDP and the G-protein binding residues for (b) unphosphorylated and (d) phosphorylated G*α*i3. The interdomain separation between Ras-like and helical domains for (c) unphosphorylated and (e) phosphorylated G*α*i3. Heatmap of GDP’s contact fraction with nucleotide binding residues for (f) unphosphorylated and (g) phosphorylated G*α*i3 for each state.

Further, we examined the domain separation in each metastable state for both systems. The unphosphorylated G*α*i3 exhibited a peak at 14 Å for the state UB and close to 17 Å for UI, indicating an opening of domains in the intermediate state that could facilitate the release of GDP from the pocket (Fig. 5c). The broader distribution observed for the U0 is clearly suggestive of increased interdomain flexibility, consistent with the NMR and MD simulation studies reported for the nucleotide-free G-proteins.(97) In the phosphorylated G*α*i3, major peak values for the states PB and PI1 appeared close to 14 Å though an increased population of large domain separation was observed for the state PI1 while transitioning from bound to this state (Fig. 5e). In fact, the marginal decrease in average GDP contacts observed for PI1 (Fig. 5d) largely originated from the helical domain as a result of its separation from the Ras-like domain. For the state PI2, the major population was observed at a value of 20 Å, whereas domain separation for the unbound P0 showed much larger values as compared to U0 (Fig. 5e). Interestingly, the reduction in average contacts of GDP in UI and PI2 are similar (Figs. 5b and 5d); however, the extent of domain separation is found to be notably different in these two intermediate states (Figs. 5c and 5e), which is suggestive of the differential interplay of domain dynamics during GDP release for the systems under study.

To elucidate the steps of the GDP dissociation in both systems, we looked into the nucleotide contacts of G protein residues for each metastable state. While transitioning from the bound state UB to the intermediate state UI, primarily the contacts of residues from the P-loop (in particular, G42^G.h1s1.3^, S44^G.h1.5^, K46^G.H1.1^, and S47^G.H1.2^) and sw1 (T177^G.hfs2.1^, R178^G.hfs2.2^, and V179^G.hfs2.3^) were observed to be weakened in the unphosphorylated G*α*i3 (Figs. 5a and 5f). A subsequent loss of contacts from P-loop, NDS motif, sw1, *β* -strand 5 (S5), *α*G, s5hg loop, and s6h5 loop resulted in the formation of the unbound state U0. On the other hand, for the phosphorylated G*α*i3, the weakening of contacts with the residues of NDS motif (N149^H.hdhe.4^, D150^H.hdhe.5^, and S151^H.HE.1^) and sw1 (T177^G.hfs2.1^ and V179^G.hfs2.3^) was found to be pivotal in transition to the intermediate state PI1 from the bound state PB (Figs. 5a and 5g). Since the NDS motif is present in the helical domain, the relatively higher weakening of its GDP contacts is primarily attributed to a substantial interdomain separation, observed more frequently for the phosphorylated than the unphosphorylated G*α*i3 as discussed previously (Figs. 5c and 5e). Whereas, for the intermediate state PI2, the major reduction in contacts was observed for the residues in P-loop (notably, G42^G.h1s1.3^, K46^G.H1.1^, and S47^G.H1.2^) and sw1 (R178^G.hfs2.2^). Moreover, additional P-loop residues (S44^G.s1h1.5^ and G45^G.s1h1.6^) also exhibited a mild weakening of contacts, thereby contributing to an overall decrease of the affinity of GDP molecule. Eventually, the fully dissociated state (P0) was attained by the loss of contacts with the rest of the Ras-like domain residues (*β* -strand 5, *α*G, s5hG loop, and s6h5 loop). Therefore, these results suggest that for the phosphorylated G*α*i3, the GDP dissociation initiates by losing its contact with the NDS motif of the helical domain and sw1 region due to the increased interdomain flexibility caused by the unfolding of *α*F helix (Fig. 3b). Further, the loss of contact with the P-loop residues, followed by other structural elements leads to the state P0. In contrast, unphosphorylated G*α*i3 undergoes an initial reduction in contacts with the P-loop, whereby a decrease in binding affinity of GDP resulted in a weak association with the sw1 residues, before finally going to the unbound state U0.

The lifetime (*τ*_*l*_) obtained for each state from the HMM model showed that the GDP-bound state of the unphosphorylated G-protein is significantly more stable than that of the phosphorylated system (*τ*_*l*_ for UB: ∼1710 *μ*s, whereas for PB: ∼0.2 *μ*s, Figs. 6a and 6b). The orders of magnitude larger lifetime obtained for the unphosphorylated G*α*i3 may in fact be correlated with the previous findings of lack of GDP release in microseconds-long simulations of Gt (a subtype of G-protein).(16) Additionally, the mean first passage time (MFPT) estimate for the transition of UB to UI is nearly three orders of magnitude larger than for UI to U0 (Fig. 6c). In contrast, for the phosphorylated G*α*i3, transitions from the bound state PB to the unbound state P0 via intermediates, PI1 and PI2, are significantly faster (MFPT *<*0.6 *μ*s, Fig. 6d). This clearly indicates that the rate of GDP dissociation increases notably due to the phosphorylation at Y154^H.HE.4^ and Y155^H.HE.5^ in G-protein.

**Figure 6:**
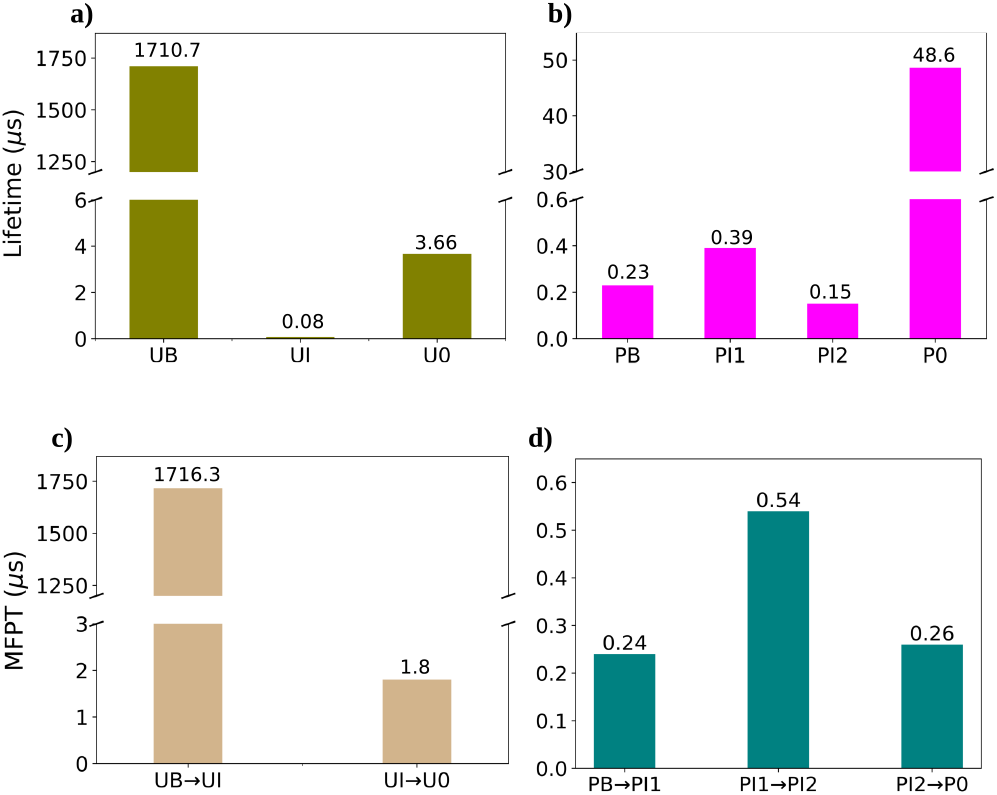
The lifetimes obtained from MSM for the states of (a) unphosphorylated and (b) phosphorylated G*α*i3. The mean first passage time (MFPT) between bound to unbound states for (a) unphosphorylated and (b) phosphorylated G-proteins.

## 4 Conclusions

The hyperactivation and dysregulation of the G-protein signaling pathways are implicated in various diseases, including diabetes, organ fibrosis, and cancer. In pathophysiological signaling, RTK-mediated phosphorylations at residues Y154^H.HE.4^ and Y155^H.HE.5^ are shown to enhance the activation of G*α*i3 by a faster exchange of GDP for GTP. To understand the mechanism of this process, we successfully employed WTMetaD and unbiased MD simulations and captured the nucleotide release in the presence and absence of phosphorylations. We showed that the phosphorylations at Y154^H.HE.4^ and Y155^H.HE.5^ in helix *α*E influences the interdomain dynamics, resulting in a considerable separation between the helical and Ras-like domains of G*α*i3. Significantly, the weakening of salt bridge interactions at the domain interface contributes to the increased distance between the domains. Moreover, helix *α*F, which is positioned opposite to helix *α*E, was unfolded in the phosphorylated system due to the disruption of interactions that stabilize its secondary structure. This increased the flexibility near the hinge region, contributing to the larger domain separation. Furthermore, the release of GDP in the phosphorylated G-protein was influenced by fluctuations in the P-loop and sw1, which directly disrupted the interactions between the nucleotide and residues of these regions. The sw2 also exhibited greater dynamics in response to phosphorylations, potentially relieving spatial constraints and thereby allowing fluctuations in neighboring structural units. A similar mechanism involving sw2 was suggested for GPCR-mediated G-protein activation in a previous study (96), indicating parallels with the phosphorylation-based activation of G*α*i3. Additionally, Markov state models revealed differences in intermediates formed during GDP release and quantified the rates of transition between the states in the unphosphorylated and phosphorylated systems, demonstrating a faster release in the latter compared to the former in consensus with previous experimental results. In summary, our study provided atomistic insights into the phosphorylation-induced activation of G-proteins and identified the specific structural motifs involved in this process. The knowledge obtained from this study could potentially facilitate the targeting of non-canonical G-protein signaling pathways and the development of therapeutic strategies to address disorders resulting from dysregulations. Nevertheless, we acknowledge that additional investigations into other G-protein subtypes are necessary to ascertain the universality of the phosphorylation-mediated GDP dissociation mechanism.

## CRediT authorship contribution statement

**K.S**.: Investigation, Formal analysis, Data curation, Visualization, Writing - original draft. **M.K.M**.: Writing - original draft. **R.K.M**.: Conceptualization, Project administration, Resources, Supervision, Funding acquisition, Writing - original draft.

## Acknowledgments

The authors acknowledge the high-performance computing (HPC) facility of IISER Bhopal. K.S. is supported by the research fellowship provided by IISER Bhopal, and M.K.M. is supported by the research fellowships provided by the Council of Scientific and Industrial Research (CSIR), India, and IISER Bhopal. R.K.M. gratefully acknowledges the financial support provided by the Science and Engineering Research Board (SERB), Department of Science and Technology, India (file no. CRG/2023/000970). The support and resources provided by PARAM Sanganak under the National Supercomputing Mission, Government of India at the Indian Institute of Technology, Kanpur are gratefully acknowledged.

## SUPPLEMENTARY MATERIAL

An online supplement to this article can be found by visiting BJ Online at http://www.biophysj.org.

