## Supplemental File for "Mechanistic Insights into G-protein Activation via Phosphorylation Mediated Non-Canonical Pathway"

### Supplementary Material for: "Mechanistic Insights into G-protein Activation via Phosphorylation Mediated Non-Canonical Pathway"

Kunal Shewani[1], Midhun K. Madhu[2], and Rajesh K. Murarka*[1]

[1]Department of Chemistry, Indian Institute of Science Education and Research Bhopal, Bhopal Bypass Road, Bhopal 462066, MP, India.
[2]Department of Biological Sciences, Indian Institute of Science Education and Research Bhopal, Bhopal Bypass Road, Bhopal 462066, MP, India.

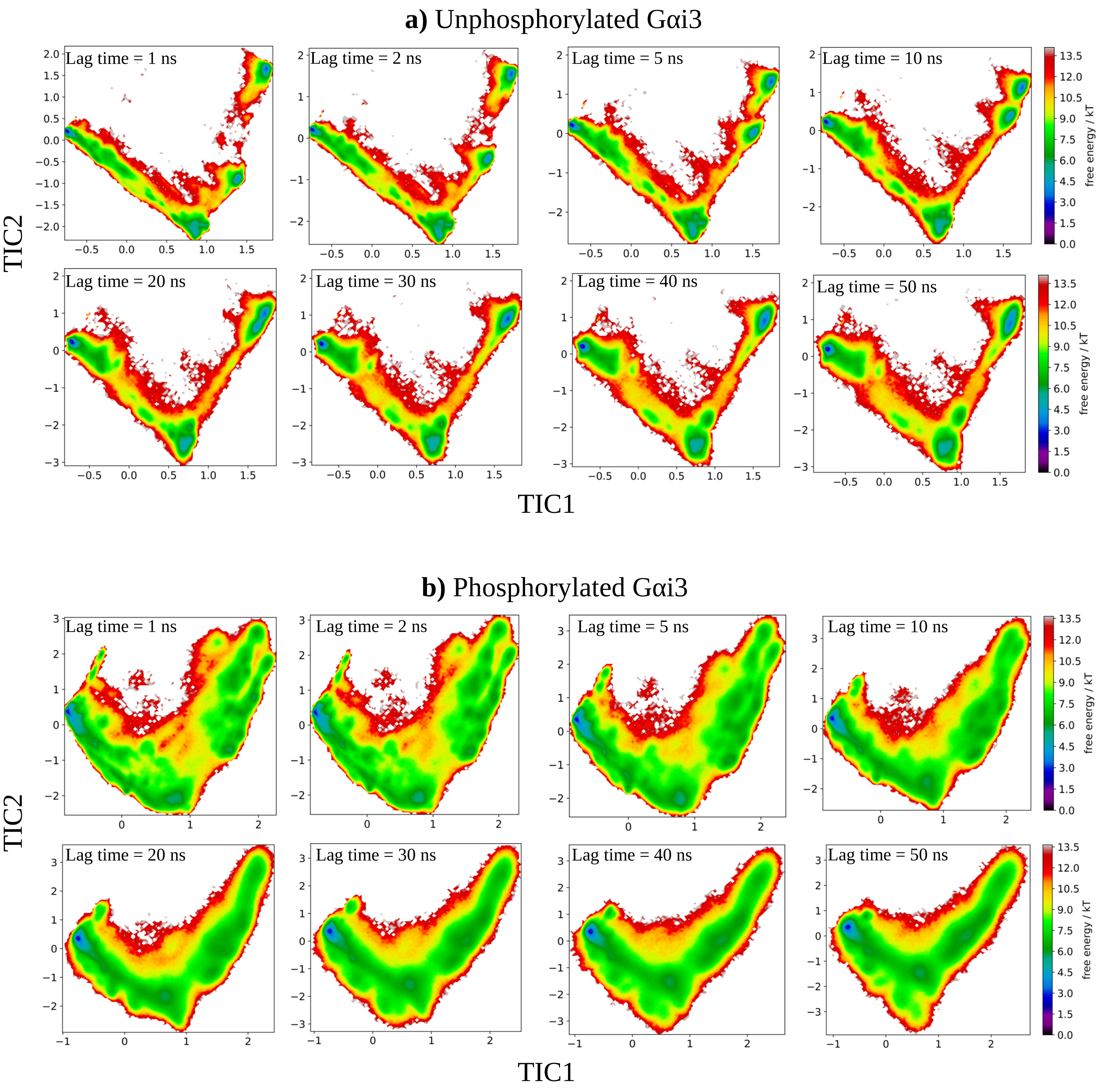


Figure S1: The tICA free energy surface, projection on the first two independent components (TICs), for (a) unphosphorylated, and (b) phosphorylated G-protein systems with varied lag times when GDP heavy atom contacts were used as input feature.

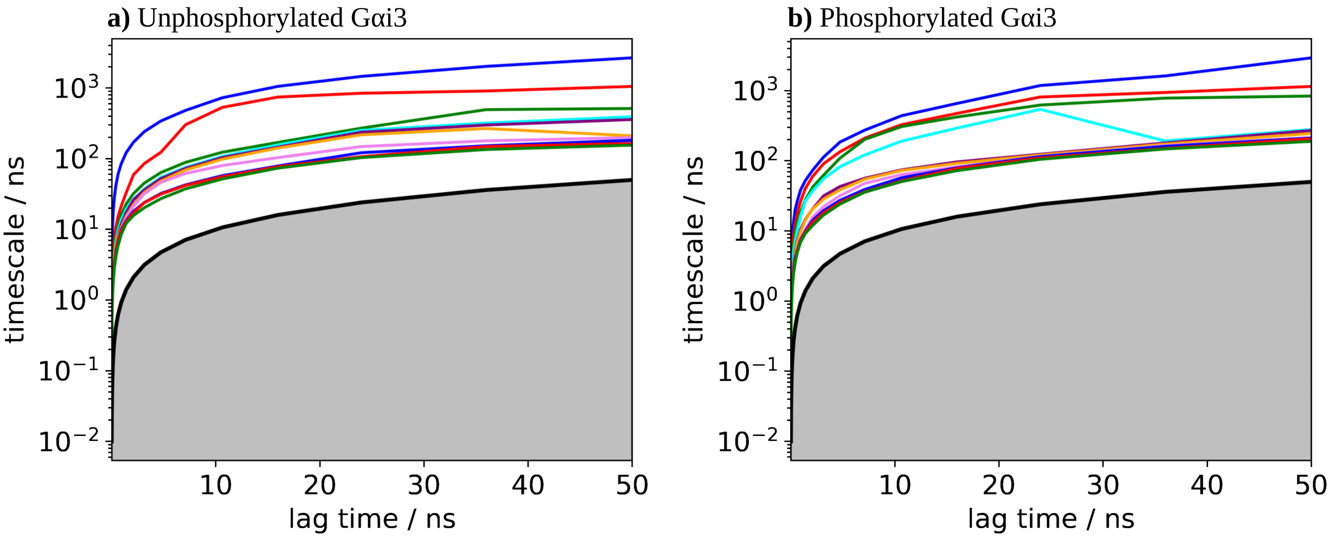


Figure S2: Implied timescales plot for (a) Unphosphorylated, and (b) Phosphorylated G$\alpha$i3.

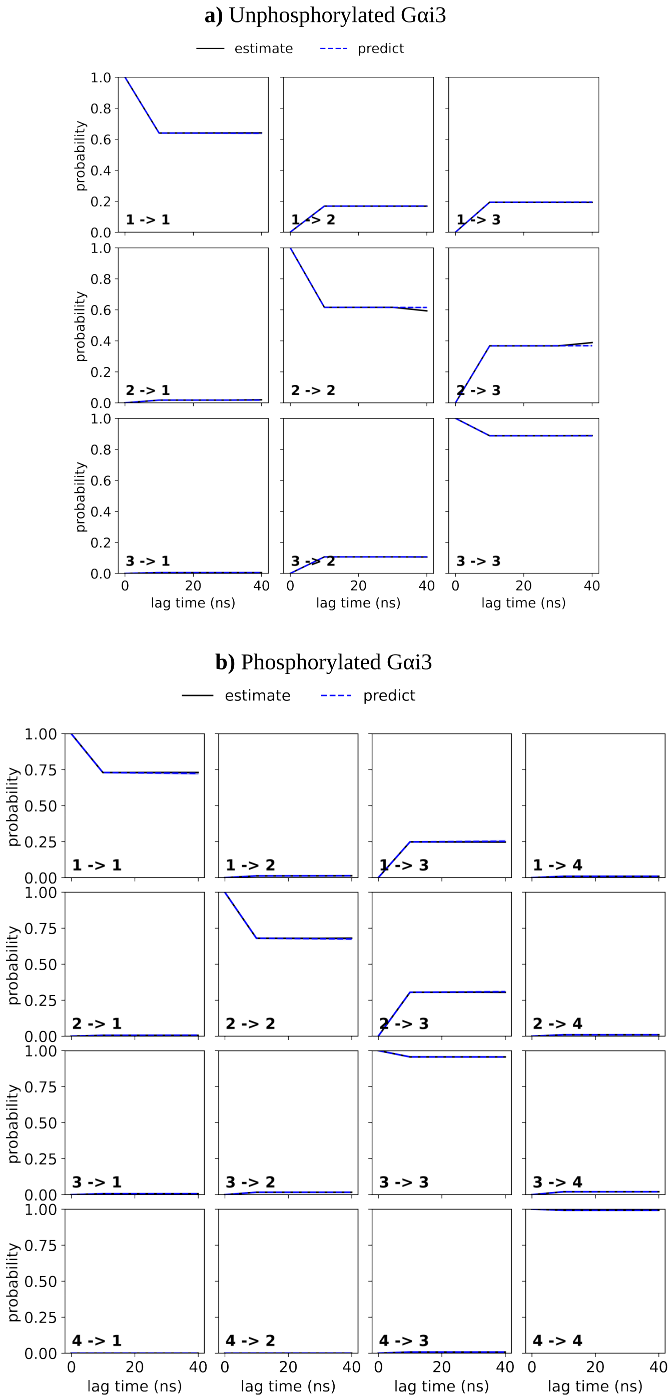


Figure S3: Chapman-Kolmogrov test for a three and four-state model of (a) unphosphorylated and (b) phosphorylated systems, respectively.

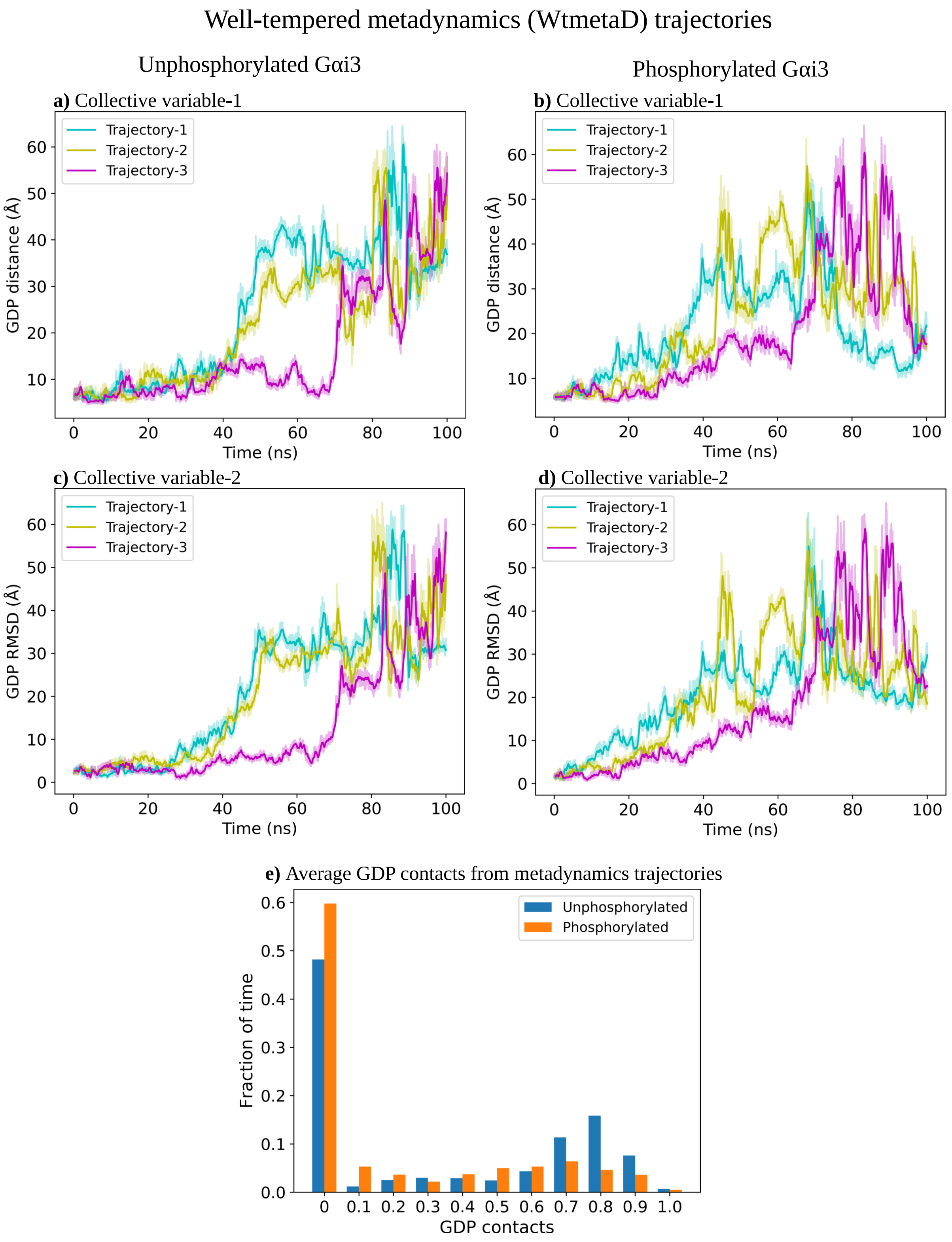


Figure S4: Timeseries of the collective variables used for well-tempered metadynamics simulations for unphosphorylated and phosphorylated G-proteins. (a) and (b) GDP's phosphate atoms distance from the backbone of residues K46$^{G.H1.1}$, S47$^{G.H1.2}$, and T48$^{G.H1.3}$. (c) and (d) GDP's RMSD to the initial bound state. (e) Average GDP's contact fraction versus fraction of time for all the trajectories.

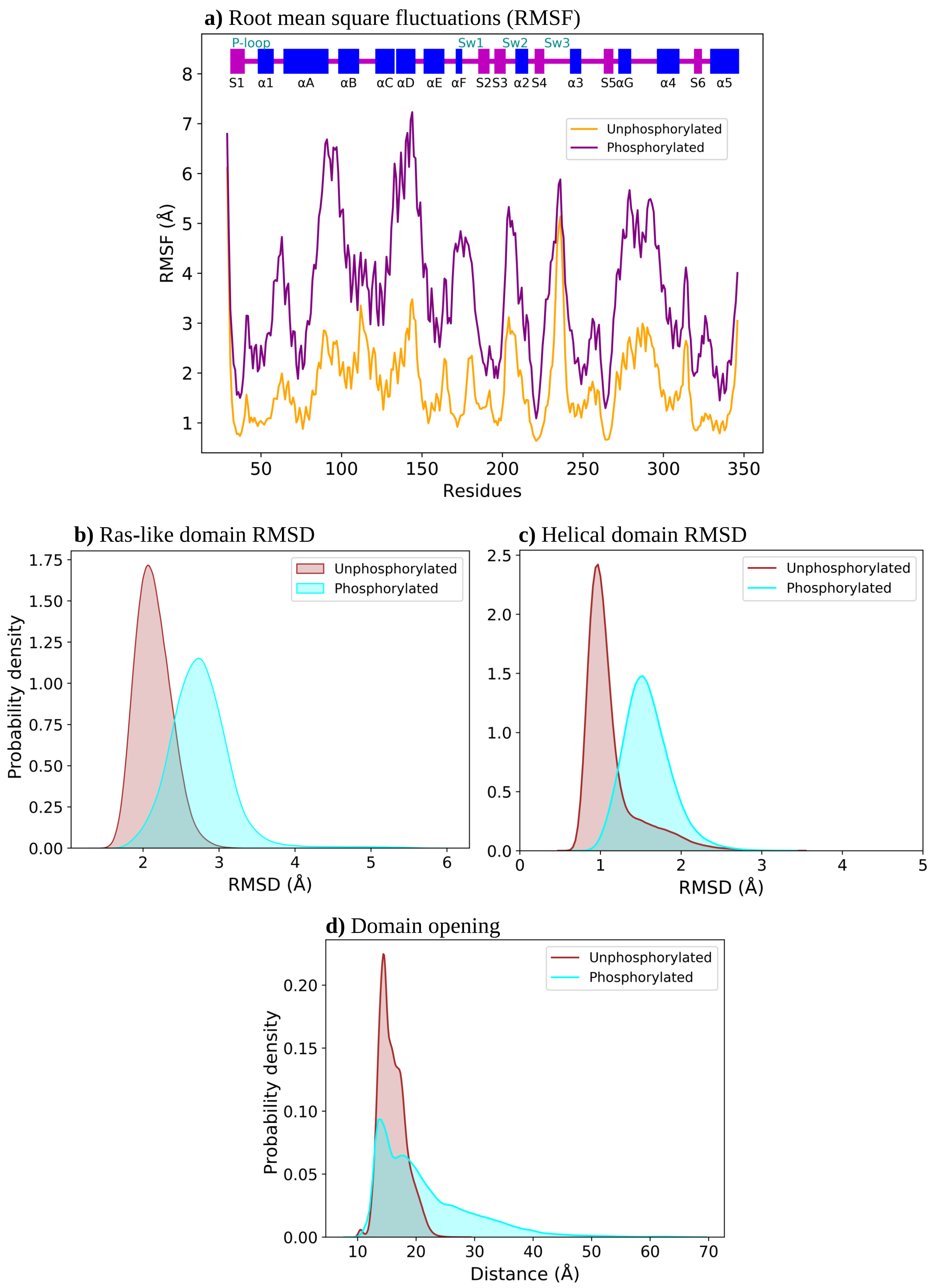


Figure S5: (a) RMSF of residues for unphosphorylated and phosphorylated G$\alpha$i3. RMSD distribution plots of (b) Ras-like domain and (c) Helical domain for both the systems. (c) Distributions of nterdomain separation for each system.

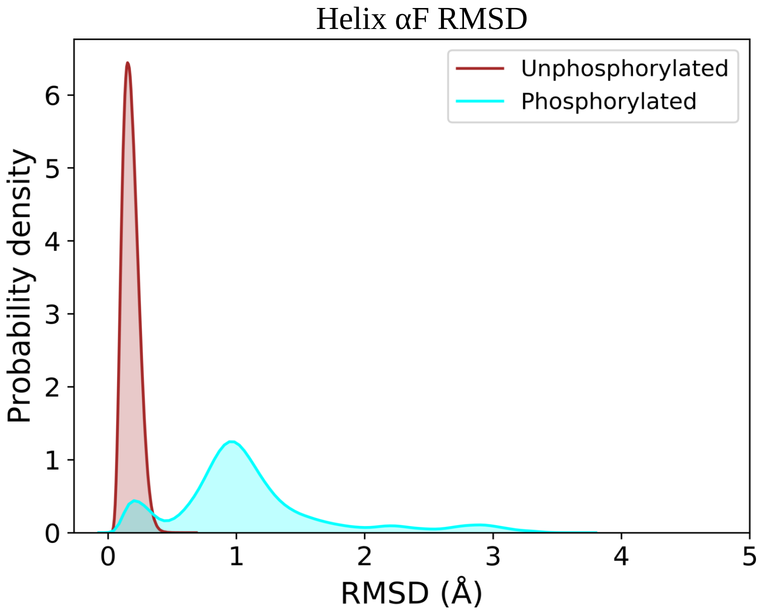


Figure S6: The RMSD distribution of helix $\alpha$F for the unphosphorylated and phosphorylated G-protein.

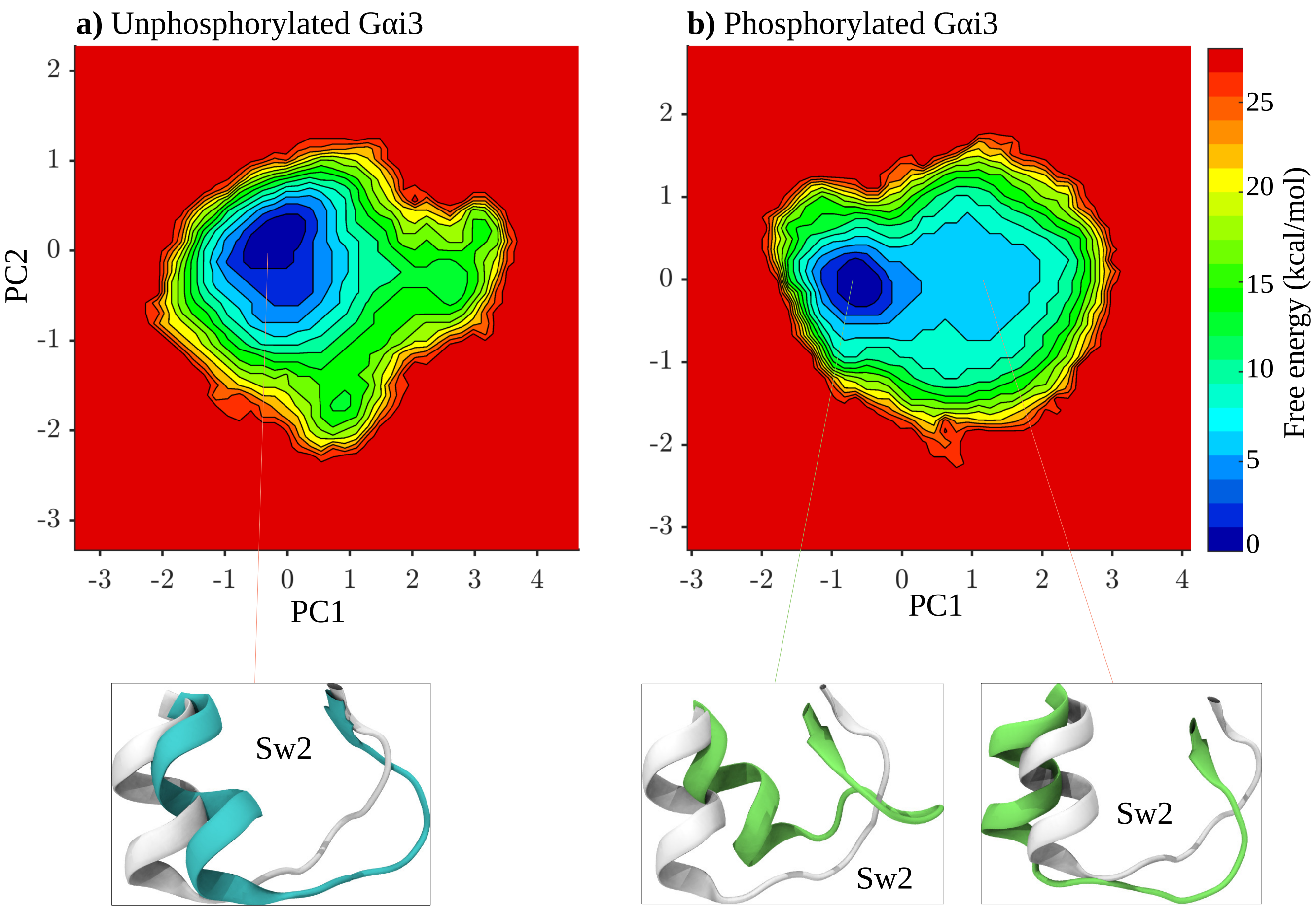


Figure S7: Principal component analysis (PCA) of $C_\alpha$ atoms of switch 2 after aligning the Ras-like domain. The free energy surface along first two principal components (PCs) (a) unphosphorylated and (b) phosphorylated G$\alpha$i3 with the representative conformations from minima.

Table S1: Average interaction energy (kcal/mol) of residue pairs at domain interface and with Y154$^{H.HE.4}$ and Y155$^{H.HE.5}$ in unphosphorylated and phosphorylated G$\alpha$i3. The standard error of the mean gives the error.

| **Residue pair** | **Unphosphorylated G$\alpha$i3** | **Phosphorylated G$\alpha$i3** |
|---|---|---|
| K54$^{G.H1.9}$–E65$^{H.HA.3}$ | -91.2 ± 1.1 | -57.6 ± 1.9 |
| K54$^{G.H1.9}$–Y154$^{H.HE.4}$ | 0.3 ± 0.003 | -31.3 ± 0.4 |
| K54$^{G.H1.9}$–Y155$^{H.HE.5}$ | 0.2 ± 0.01 | -40.8 ± 0.4 |
| E65$^{H.HA.3}$–Y154$^{H.HE.4}$ | -0.2 ± 0.01 | 41.8 ± 0.2 |
| E65$^{H.HA.3}$–Y155$^{H.HE.5}$ | -0.2 ± 0.01 | 41.8 ± 0.2 |
| R144$^{H.HD.11}$–D229$^{G.s4h3.3}$ | -34.1 ± 1.1 | -25 ± 0.7 |
| R144$^{H.HD.11}$–Y154$^{H.HE.4}$ | 0.5 ± 0.02 | -37.3 ± 0.8 |
| R144$^{H.HD.11}$–Y155$^{H.HE.5}$ | 0.4 ± 0.01 | -32.8 ± 0.3 |
| D229$^{G.s4h3.3}$–Y154$^{H.HE.4}$ | -0.9 ± 0.01 | 37.9 ± 0.6 |
| D229$^{G.s4h3.3}$–Y155$^{H.HE.5}$ | -0.4 ± 0.01 | 38.1 ± 0.5 |
| R144$^{H.HD.11}$–D231$^{G.s4h3.5}$ | -51.9 ± 1.3 | -39.6 ± 1.6 |
| D231$^{G.s4h3.5}$–Y154$^{H.HE.4}$ | -0.6 ± 0.01 | 31.2 ± 0.4 |
| D231$^{G.s4h3.5}$–Y155$^{H.HE.5}$ | -0.3 ± 0.01 | 28.8 ± 0.3 |
| D150$^{H.hdhe.5}$–K270$^{G.s5hg.1}$ | -86.4 ± 1.3 | -46.4 ± 2.1 |
| D150$^{H.hdhe.5}$–Y154$^{H.HE.4}$ | -3.2 ± 0.1 | 53.9 ± 0.8 |
| D150$^{H.hdhe.5}$–Y155$^{H.HE.5}$ | -0.9 ± 0.02 | 50.9 ± 0.4 |
| K270$^{G.s5hg.1}$–Y154$^{H.HE.4}$ | 1.3 ± 0.01 | -48.8 ± 1.2 |
| K270$^{G.s5hg.1}$–Y155$^{H.HE.5}$ | 0.4 ± 0.01 | -47.7 ± 0.7 |
| E145$^{H.HD.12}$–R280$^{G.HG.12}$ | -24.7 ± 5 | -16.6 ± 4.8 |
| E145$^{H.HD.12}$–Y154$^{H.HE.4}$ | -0.2 ± 0.1 | 25.5 ± 2.7 |
| E145$^{H.HD.12}$–Y155$^{H.HE.5}$ | -0.3 ± 0.1 | 24.7 ± 1.2 |
| R280$^{G.HG.12}$–Y154$^{H.HE.4}$ | 0.3 ± 0.2 | -39.5 ± 16.3 |
| R280$^{G.HG.12}$–Y155$^{H.HE.5}$ | 0.3 ± 0.1 | -29.3 ± 4.1 |
| D150$^{H.hdhe.5}$–K277$^{G.HG.12}$ | -42 ± 12 | -34.1 ± 13.8 |
| K277$^{G.HG.12}$–Y154$^{H.HE.4}$ | 0.7 ± 0.1 | -36.2 ± 7.6 |
| K277$^{G.HG.12}$–Y155$^{H.HE.5}$ | 0.3 ± 0.1 | -32.1 ± 4.6 |
